# Caste-Specific Proboscis Extension Responses in Honey Bees to Sucrose and Royal Jelly Stimuli

**DOI:** 10.64898/2026.01.23.701277

**Authors:** Babur Erdem, Sedat Sevin, Okan Can Arslan, Ayse Gul Gozen, Hande Alemdar, Ali Emre Turgut, Tugrul Giray, Erol Sahin

## Abstract

Understanding the nutritional preferences of honey bees (*Apis mellifera*) is essential for comprehending their behavioral ecology and the division of labor within a colony. While gustatory sensitivity to sucrose is well-documented in workers, a significant research gap exists regarding the sensory responses of queens and their reactions to caste-specific nutrition such as royal jelly. This study utilized the proboscis extension response (PER) assay to compare the food preferences of three distinct bee categories: foragers, 1-day-old workers, and queens. Subjects were presented repeatedly, in a pseudorandom order, with water, sucrose, royal jelly, and a sucrose–royal jelly mixture as gustatory stimuli. Foragers exhibited a high responsiveness to sucrose and showed uniformly low responsiveness to other stimuli. Although 1-day-old workers showed high responsiveness to sucrose, unlike foragers, they also responded to the sucrose–royal jelly mixture. Queens displayed a unique response profile, with near-ceiling responsiveness to both royal jelly and the mixture, followed by response to sucrose solution without habituation. Additionally, responsiveness to the sucrose was higher in foragers than in 1-day-old workers. These findings suggest that the honey bee gustatory and sensory system is tuned to the specific nutritional requirements of caste and age.

## 1. INTRODUCTION

Understanding the food preferences of honey bees (*Apis mellifera*) is crucial for unraveling the behavioral ecology of this key pollinator species. Food selection not only affects individual bee survival and colony-level resource allocation but also influences feeding dynamics, trophallactic interactions, and division of labor (Farina & Núñez, 1991; de Brito Sanchez, 2011). These preferences are driven by physiological requirements, sensory response thresholds, and the specific tasks the bees perform.

The food sources of honey bees vary by their caste (i.e., queens versus workers), and their temporal (i.e., foragers versus nurses) and behavioral division of labor (i.e., pollen versus nectar foragers). Temporal polyethism is an age-related division of labor where younger bees carry out in-hive tasks such as nursing, while older bees take on outside jobs such as nectar and pollen foragers (Seeley, 1982). Pollen is the primary natural source of protein, and nurses consume 3.4 to 4.3 mg pollen per day (Crailsheim et al, 1992). Pollen provides essential amino acids, including arginine, histidine, lysine, tryptophan, phenylalanine, methionine, threonine, leucine, isoleucine, and valine (de Groot, 1952). This protein source is essential for the development of their hypopharyngeal glands, to synthesize the protein-rich jelly used to feed larvae and the queen (Crailsheim, 1990a). In addition to pollen consumption, young bees receive jelly via trophallaxis from older nurse bees (Crailsheim, 1990a; Free, 1957; Crailsheim, 1990b). Nurses also receive nectar collected by foragers. However, the sugar concentration in the crops of 5 to 6-day-old worker bees is typically lower than that found in the crops of active foragers (Pankiw et al., 2004). Foragers primarily consume carbohydrates to fuel the intense metabolic cost of flight. The transition to foraging causes physiological shifts. Hypopharyngeal gland atrophy occurs, and proteolytic enzyme activity decreases significantly in foragers (Crailsheim & Stolberg, 1989). Especially, pollen consumption decreases with age (Lotmar, 1938; Lindauer, 1952). Thus, the diet shifts from protein accumulation to carbohydrate catabolism (Crailsheim, 1990a). The queen is fed royal jelly. The queen receives protein that has already been processed and synthesized by the nurse bees (Crailsheim, 1990a). This high-protein diet supports her highly developed reproductive characteristics (Fèvre & Dearden, 2024).

The proboscis extension response (PER) is widely considered a standard and established assay for quantifying both gustatory responsiveness and appetitive learning (Scheiner et al., 2003). The PER can be utilized in a non-associative learning assay, which is especially used to test gustatory responsiveness or non-associative forms of learning. This is measured by repeatedly stimulating the antennae with a solution and observing the frequency or magnitude of the proboscis extension over time (Scheiner et al., 2003). Honey bees give PER to sucrose when the solution comes into contact with their antennae (Bitterman et al. 1983). Newly emerged bees typically have low responsiveness to sucrose, and when they approach foraging age, their responsiveness increases (Pankiw & Page, 1999). Sucrose responsiveness is also higher in foragers than in nurse bees and is related to the expression of specific genes in the brain (Degirmenci et al., 2018). In addition, young bees exhibit a higher habituation rate to sucrose than older bees (Guez et al., 2001; Scheiner et al., 2003). Thus, sucrose responsiveness generally increases as bees age; younger bees are less responsive to sucrose. Another factor that influences responsiveness is the social context: brood pheromone lowers the sucrose response thresholds of young bees (Pankiw & Page, 2001). Foragers also vary in sucrose response. The sucrose sensitivity of foragers correlates with their foraging specialization. Water and pollen foragers have the highest responsiveness to sucrose, followed by nectar foragers (Scheiner et al., 2001). Queen bees are also responsive to sucrose (Gong et al., 2018). In addition to sucrose, worker bees also give PER to pollen (Nicholls & Hempel de Ibarra, 2013), and pollen foragers are more responsive to pollen (Moreno & Arenas, 2023; Moreno & Arenas, 2024).

While taste sensitivity to sucrose and PER-based learning in honey bees have been extensively characterized, particularly across age groups and nutritional specializations, there is a gap regarding caste-comparative PER studies involving queens and queen-related nutritional stimuli (royal jelly). Based on the caste-specific nutritional source of honey bees and the queen’s royal jelly-based diet, we hypothesize that PER will differ for caste-specific stimuli. We predict foragers and young workers will show the highest sensitivity to sucrose, while queens will show high sensitivity to royal jelly. We test this hypothesis using a PER assay in which foragers, 1-day-old workers, and queens are presented with water, sucrose, royal jelly, and a royal jelly– sucrose mixture across repeated, pseudorandomized trials.

## 2. METHOD

### 2.1 Collecting bees and handling

Proboscis extension response (PER) experiments were conducted with three bee categories: foragers, 1-day-old workers, and queens. Returning foragers were collected at the hive entrance using a plastic mesh (Giray et al., 2007) and placed in flight boxes, where they were food-deprived for 2 h. For the 1-day-old group, capped brood combs were removed from the hives and placed in a box inside an incubator at 35 °C and 65% relative humidity overnight. Newly emerged 1-day-old workers were collected 2 h before the experiment and placed in flight boxes for food deprivation. Both foragers and 1-day-old workers were obtained from two source colonies. After the food-deprivation period, bees were harnessed and kept ready for testing.

Queen bees were obtained from three different beekeepers. Mated queens were collected from their mini-nucs, small colonies with ∼2000 workers, where each queen was laying eggs, and was attended by her worker offspring. Each queen was transferred to our lab in a queen cage with 5– 10 attendant nurse bees and placed in the incubator until the experiment (Fahrbach et al. 1995). Before testing, queens were separated from the nurses, harnessed, and prepared for PER assays as described for the worker groups.

### 2.2 Test procedure

Four test solutions were prepared: water, pure royal jelly, 50% sucrose solution (w/v), and a mixture consisting of 50% royal jelly, 25% sucrose, and 25% water (w/w/v). To assess PER, separate cotton swabs were dipped into each solution and brought into contact with the antennae of the bee for 3 s. Each bee received a total of 60 trials presented in a pseudorandom order (S, W, RJ, M, S, M, W, RJ, M, S, W, RJ, M, W, S, RJ, M, W, RJ, S, W, S, RJ, M, S, M, W, RJ, M, S, W, RJ, M, W, S, RJ, M, W, RJ, S, W, S, RJ, M, S, M, W, RJ, M, S, W, RJ, M, W, S, RJ, M, W, RJ, S), where S = sucrose, W = water, RJ = royal jelly, and M = mixture, each stimulus appears 15 times overall with no consecutive repeats. The sequence prevents conditioning to a visual cue, such as an upcoming cotton swab, and creates a similar sensitization or habituation effect for all solutions. The inter-trial interval was 1 min for each bee. For each stimulation, the presence (1) or absence (0) of PER was recorded. The forager and 1-day-old worker groups each consisted of 24 individuals, and the queen group consisted of 10 individuals.

### 2.3 Statistical analysis

All statistical analyses were conducted in RStudio using the R version 4.3 (R Core Team, 2020). Data manipulation and reshaping were performed using the *readxl, dply*r, *tidyr*, and *stringr* packages.

While raw PER ratios (the total number of positive responses divided by 15 trials per bee) were calculated to provide a descriptive overview of the data, these summary statistics do not account for individual variation or temporal effects like habituation. Therefore, formal inference was based on a binomial generalized linear mixed model (GLMM) with a logit link.

Because PER is a binary outcome (0 or 1) recorded repeatedly from the same individuals across trials and stimuli, inference was based on a GLMM rather than independent-samples tests. A binomial GLMM with a logit link was fitted using the g*lmer* function in the *lme4* package, including bee categories (forager, 1-day-old, queen) and stimulus (water, sucrose, royal jelly, mixture) to test whether stimulus responsiveness differed among bee categories. The repeated-measures structure was accounted for by including bee identity as a random intercept, allowing baseline response probability to vary among individuals. This modeling choice is appropriate because it respects the non-independence of trials within bees, uses the correct distribution for binary outcomes, and yields interpretable bee categories-by-stimulus effects while retaining trial-level information. We therefore analyzed PER using binomial logistic models, which estimate effects on the log-odds scale and then convert them to predicted response probabilities via the logistic link. Using probabilities provides model-based estimates that account for the discrete nature of the data and the non-independence introduced by repeated measures within bees.

To capture systematic changes in responsiveness across repeated stimulation (e.g., habituation or sensitization), trial order was included as a scaled linear term (trial) and a quadratic term (trial^2^). Both trial-order terms were retained in the final model.

Model adequacy was evaluated using diagnostic functions from the *performance* package, including checks for overdispersion and singularity. Predicted probabilities were obtained as estimated marginal means (EMMs) on the response scale using the *emmeans* package, which back-transforms log-odds to PER probabilities and provides confidence intervals. Multiple comparisons among EMMs were conducted as Tukey-adjusted pairwise contrasts.

To test habituation within each stimulus, we evaluated whether PER decreased across the 15 repeated presentations of that stimulus. We fitted a binomial generalized linear mixed model (GLMM) with a logit link, using the within-stimulus repetition index (1–15) as the focal predictor and including bee identity as a random intercept to account for repeated measures within individuals. The repetition index was z-standardized prior to model fitting to improve numerical stability and allow comparable effect scaling across groups. Model fitting used maximum likelihood with the Laplace approximation. Habituation was inferred when the repetition coefficient was negative and statistically different from zero (α = 0.05). Inference was based on a one-sided test derived from the Wald *z* statistic, and p-values were adjusted across the set of caste × stimulus tests using the Holm procedure to control the family-wise error rate.

## 3. RESULTS

### 3.1 PER ratios to gustatory stimuli

Foragers responded most strongly to sucrose, with comparatively low and similar responses to royal jelly, the mixture, and water. In 1-day-old workers, response to sucrose was again the highest, and the mixture elicited responses comparable to sucrose, whereas royal jelly and water elicited lower responses. In contrast, queens showed near-ceiling PER to royal jelly and the mixture, both stimuli exceeding sucrose and water. Foragers (*n* = 24) exhibited a high mean and median PER ratio for sucrose (*μ* = 0.931, *M* = 1) but low ratios for water (*μ* = 0.403, *M* = 0.467), royal jelly (*μ* = 0.386, *M* = 0.400), and the mixture (*μ* = 0.411, *M* = 0.400). In 1-day-old workers (*n* = 24), sucrose elicited the strongest response (*μ* = 0.792, *M* = 0.867), followed by the mixture (*μ* = 0.619, *M* = 0.667), royal jelly (*μ* = 0.494, *M* = 0.467), and water (*μ* = 0.439, *M* = 0.467). Queens (*n* = 10) exhibited near-maximal responsiveness to both the mixture (*μ* = 0.993, *M* = 1) and pure royal jelly (*μ* = 0.993, *M* = 1). In contrast, sucrose yielded lower responses (*μ* = 0.800, *M* = 0.900), and water elicited the minimum response ratio (*μ* = 0.160, *M* = 0.100).

We then analyzed the proboscis extension responses (PER: 1, no PER: 0) (Figure 1A-C) at the trial level using a binomial generalized linear mixed model (GLMM) with a logit link, with stimulus (sucrose, water, royal jelly, and mixture), bee categories (forager, 1-day-old worker, queen), and their interaction as fixed effects and a random intercept for individual bee. The model indicated strong effects of trial order in scaled trial and its square (standardized trial: *z* = −18.39, *p* < 0.001; trial^2^: *z* = 5.13, *p* < 0.001). Model diagnostics indicated no overdispersion and non-singular random effects. Estimated marginal means (EMMs) were reported as predicted probabilities with 95% confidence intervals (Table 1).

**Table 1.** Descriptive and model-based summary statistics for PER by bee categories and stimuli. For each category and stimulus condition, the table reports observed PER ratios and GLMM-derived predicted PER probabilities. PER ratio is computed as the total of positive PER numbers divided by 15 trials for each stimulus per bee. It is summarized across individuals using sample size (n), mean, standard deviation, standard error, median, and the 1st and 3rd quartiles. The GLMM columns report the estimated marginal mean PER probability on the response scale (prob), its 95% confidence interval (CI lower, CI upper), and the model-based standard error (SE model), derived from a binomial GLMM that accounts for repeated trial-level observations within individuals via a random intercept for bee identity.

| Category | Stimuli | Descriptive Statistics for PER Ratios |  |  |  |  |  |  | GLMM |  |  |  |
| --- | --- | --- | --- | --- | --- | --- | --- | --- | --- | --- | --- | --- |
|  |  | Sample Size | Mean | Standard deviation | Standard Error | Median | 1<sup>st</sup> Quartile | 3<sup>rd</sup> Quartile | prob | CI lower | CI upper | SE model |
| Forager | Mixture | 24 | 0.411 | 0.190 | 0.039 | 0.400 | 0.267 | 0.533 | 0.375 | 0.246 | 0.523 | 0.072 |
|  | Royal Jelly | 24 | 0.386 | 0.206 | 0.042 | 0.400 | 0.250 | 0.533 | 0.364 | 0.238 | 0.513 | 0.072 |
|  | Sucrose | 24 | 0.931 | 0.112 | 0.023 | 1.000 | 0.867 | 1.000 | 0.965 | 0.931 | 0.983 | 0.012 |
|  | Water | 24 | 0.403 | 0.183 | 0.037 | 0.467 | 0.317 | 0.483 | 0.369 | 0.242 | 0.517 | 0.072 |
| 1-day-old | Mixture | 24 | 0.619 | 0.269 | 0.055 | 0.667 | 0.467 | 0.817 | 0.687 | 0.542 | 0.803 | 0.068 |
|  | Royal Jelly | 24 | 0.494 | 0.276 | 0.056 | 0.467 | 0.317 | 0.683 | 0.521 | 0.371 | 0.667 | 0.078 |
|  | Sucrose | 24 | 0.792 | 0.241 | 0.049 | 0.867 | 0.600 | 1.000 | 0.887 | 0.806 | 0.937 | 0.033 |
|  | Water | 24 | 0.439 | 0.296 | 0.060 | 0.467 | 0.233 | 0.600 | 0.415 | 0.278 | 0.567 | 0.076 |
| Queen | Mixture | 10 | 0.993 | 0.021 | 0.007 | 1.000 | 1.000 | 1.000 | 0.999 | 0.990 | 1.000 | 0.001 |
|  | Royal Jelly | 10 | 0.993 | 0.021 | 0.007 | 1.000 | 1.000 | 1.000 | 0.999 | 0.991 | 1.000 | 0.001 |
|  | Sucrose | 10 | 0.800 | 0.288 | 0.091 | 0.900 | 0.867 | 0.983 | 0.907 | 0.774 | 0.965 | 0.045 |
|  | Water | 10 | 0.160 | 0.170 | 0.054 | 0.100 | 0.017 | 0.250 | 0.064 | 0.023 | 0.163 | 0.032 |

**Figure 1.**
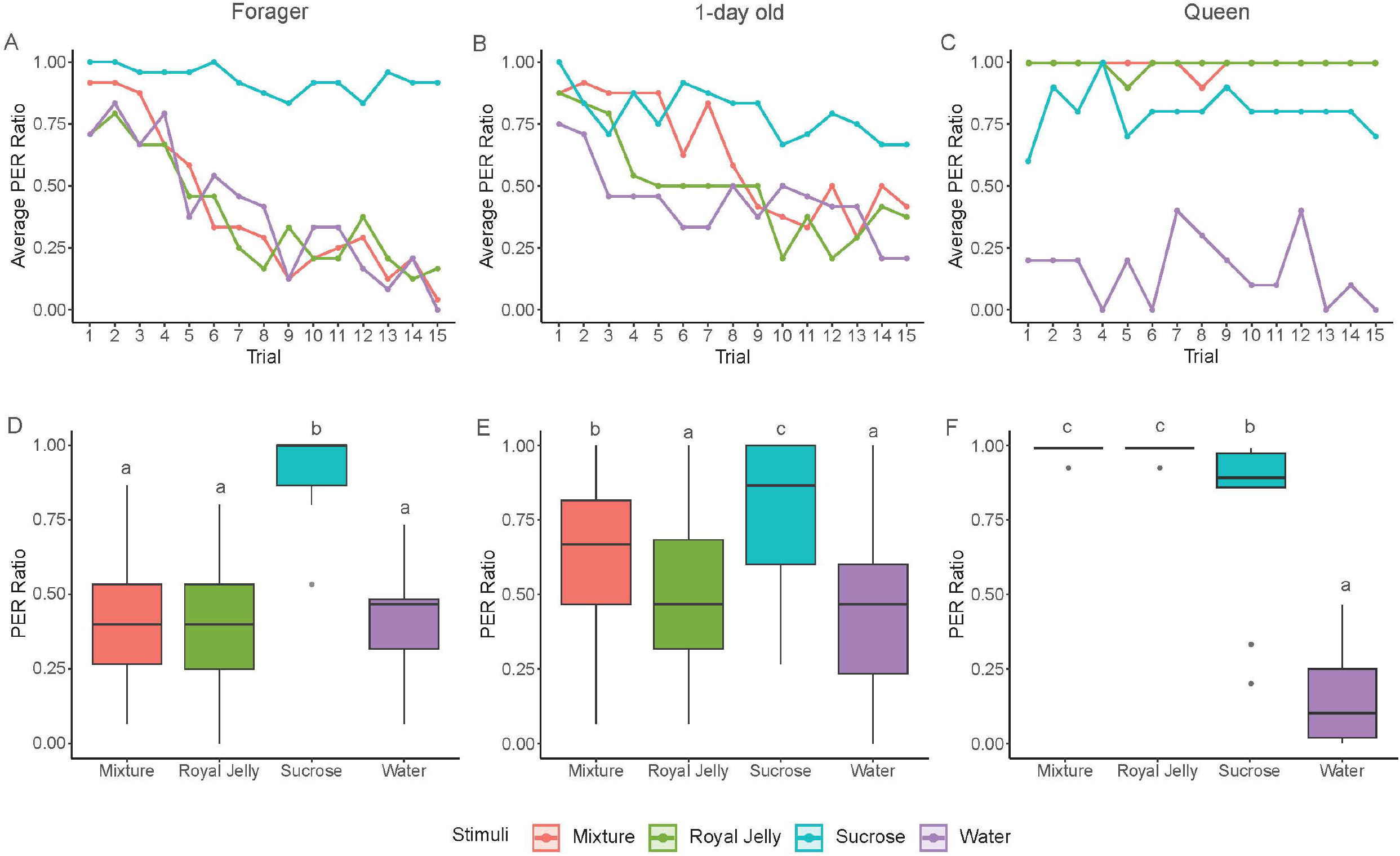
Proboscis extension response (PER) to four stimuli across honey bee categories. Panels A–C show trial-by-trial mean PER (number of positive PER/number of bees) for each stimulus in foragers (A), 1-day-old workers (B), and queens (C). Panels D–F show boxplots of the ratio of PER, which is computed as the total of positive PER numbers divided by 15 trials for each stimulus, as for sucrose, water, royal jelly, and a sucrose–royal jelly mixture in foragers (D), 1-day-old workers (E), and queens (F). Boxes are centered on the median with interquartile range, whiskers extend to 1.5×IQR. Letters above boxes are compact-letter groupings derived from the GLMM (estimated marginal means on the response scale) and indicate Tukey-adjusted pairwise differences among stimuli within each bee category, and shared letters indicate not significantly different.

### 3.2 Response patterns of bee categories

Foragers showed a strong response to the sucrose stimulus. The predicted PER probability (prob) for sucrose was high (prob = 0.965, 95% CI [0.931, 0.983]), whereas water (prob = 0.369, [0.242, 0.517]), royal jelly (prob = 0.364, [0.238, 0.513]), and mixture (prob = 0.375, [0.246, 0.523]) were uniformly low (Table 1, Figure S1).

In 1-day-old workers, PER also peaked for sucrose but showed a graded pattern across stimuli. Predicted PER was highest for sucrose (prob = 0.887, 95% CI [0.806, 0.937]), followed by mixture (prob = 0.687, [0.542, 0.803]), then royal jelly (prob = 0.521, [0.371, 0.668]) and water (prob = 0.415, [0.278, 0.567]) (Table 1, Figure S1).

Queens showed a qualitatively different profile. Predicted PER was near maximal for royal jelly (prob = 0.999, 95% CI [0.991, 1.000]) and mixture (prob = 0.999, [0.990, 1.000]), high for sucrose (prob = 0.907, [0.774, 0.965]), and low for water (prob = 0.064, [0.023, 0.163]) (Table 1, Figure S1).

### 3.3 Comparison of responses to stimuli within bee categories

Tukey-adjusted pairwise comparisons confirmed these patterns. In foragers, sucrose exceeded water, royal jelly, and mixture (all *p* < 0.001), whereas water, royal jelly, and mixture did not differ (all *p* ≥ 0.995) (Figure 1D, Table S1).

In 1-day-old workers, sucrose exceeded all other stimuli (all *p* < 0.001). The mixture also exceeded water (*p* < 0.001) and royal jelly (*p* = 0.001), while the difference between water and royal jelly was not statistically significant (*p* = 0.105) (Figure 1E, Table S1).

In queens, water was lower than each nutritive stimulus (all *p* < 0.001). Sucrose was lower than royal jelly (*p* < 0.001) and mixture (*p* < 0.001), and royal jelly and mixture did not differ (*p* = 1.000) (Figure 1F, Table S1).

### 3.4 Comparison of the responses of bee categories within each stimulus type

Bee categories differences depended strongly on the stimuli, which included water, sucrose, royal jelly, and a mixture. Queens responded less than both 1-day-old bees (*p* < 0.001) and foragers (*p* = 0.001), whereas foragers and 1-day-old workers did not differ (*p* = 0.897) (Table S2) for the water stimulus.

With sucrose as stimulus, foragers responded more strongly than 1-day-old workers (*p* = 0.027), while neither foragers (*p* = 0.236) nor 1-day-old workers (*p* = 0.938) differed significantly from queens (Table S2).

Exposure of royal jelly and mixture stimuli, queens’ PER ratio exceeded both worker groups (all *p* < .001). However, no difference was found between 1-day-old workers and foragers (*p* = 0.312) for royal jelly (Table S2).

In addition, the 1-day-old workers’ PER ratio exceeded that of foragers for the mixture stimulus (*p* = 0.009) (Table S2).

### 3.5 Habituation evaluation

Habituation was assessed as a decrease in PER probability across the 15 repeated presentations of each stimulus (Figure 1A-C) using stimulus-specific binomial mixed-effects models and defined with logit slopes (*b*), test statistic (*z*), and Holm-adjusted p values.

In foragers, PER decreased significantly across trials for water (*b* = −1.31, *z* = −8.27, *p* < 0.0001), sucrose (*b* = −0.572, *z* = −2.36, *p* = 0.046), royal jelly (*b* = −1.08, *z* = −7.31, *p* < 0.0001), and mixture (*b* = −1.67, *z* = −8.90, *p* < 0.0001) (Figure 1A). Here, the slope for sucrose is less pronounced than for other stimuli, and the significance level of the sucrose is below but close to the threshold (0.05 > *p* = 0.046)

In 1-day-old workers, PER decreased significantly across trials for all stimuli: water (*b* = −0.653, *z* = −4.71, *p* < 0.0001), sucrose (*b* = −0.587, *z* = −3.47, *p* = 0.002), royal jelly (*b* = −1.03, *z* = −6.78, *p* < 0.0001), and mixture (*b* = −1.45, *z* = −7.63, *p* < 0.0001) (Figure 1B). The slope for sucrose is also less pronounced than for other stimuli.

Here, we observed that both foragers and 1-day-old workers showed evidence of habituation for all stimuli, but the magnitude differed. Foragers exhibited a more pronounced decline in response to water, royal jelly, and mixture stimuli compared to 1-day-old workers. However, the PER ratio of foragers declined modestly (PER ratios from 1.00 on the first trial to 0.916 by the 15th trial), with a weaker negative slope for sucrose, whereas 1-day-old workers showed a sharper decline (PER ratios from 1.00 at the first trial to 0.667 at the 15th trial) with a stronger negative slope.

In queens, there was no statistically significant decrease across trials for water (*b* = −0.231, *z* = −0.943, *p* = 0.692) or sucrose (*b* = −0.0489, *z* = −0.184, *p* = 1.00). Royal jelly (*b* = 0.783, *z* = 0.659, *p* = 1.00) and the mixture (*b* = 0.236, *z* = 0.231, *p* = 1.00) also showed no statistical evidence of habituation in queens (Figure 1C). Notably, queen responses to royal jelly and mixture were near-ceiling (mean PER = 0.993), which limits sensitivity to detect decreases over trials.

## 4. DISCUSSION

This study demonstrates caste and age-related differences in PER to gustatory stimuli. A higher PER and the absence of habituation (reduction in responsiveness to repeated stimulation) for gustatory stimuli indicate a strong food preference. In the case of queen bees, these data indicate a queen-specific tuning toward royal-jelly-based stimuli. We conclude that queen bees have a very high preference for royal jelly. In contrast, foragers have a low preference for royal jelly and a high preference for high-concentration sucrose reward. On the other hand, 1-day-old bees are intermediate, with a high preference for sucrose and a lesser, but still higher, response to a royal jelly mixture than foragers. These preferences are consistent with the known feeding habits of these three bee categories, which are discussed below.

Sugar content of the royal jelly is on average 12% (w/w) (Ramanathan et al., 2011; Kolayli et al., 2025). Thus, the sugar concentration of our mixture solution should be higher than 30%, and if this percentage of sugar is not present in the royal jelly mixture, it is used as a high stimulant that breaks the habituation (Scheiner et al., 2004). However, the level of response to the mixture foragers remains at the same level as the response to the water control. One characteristic of royal jelly is its acidity, with a pH range between 3.6 and 4.2 (Ramanathan et al., 2011), which may act as a gustatory deterrent and reduce the response to sugar for foragers. Additionally, other components of royal jelly may alter feeding behavior. For instance, the addition of amino acids, such as isoleucine, proline, phenylalanine, and methionine in sucrose solution decreases its consumption rates by bees (Simcock et al., 2014). Also, proline, phenylalanine, and methionine addition to the sucrose stimuli, reduce the responsiveness (Simcock et al., 2014; Nicholls et al., 2019). These amino acids are present in royal jelly (Howe et al., 1985; Bayram et al., 2021), which may cause the reduction of PER to the mixture and royal jelly stimuli. Moreover, typically, weaker stimuli may not even cause dishabituation, but high or strong stimuli do cause dishabituation (Rankin et al., 2009). We observed habituation for mixture and royal jelly in foragers and 1-day-old workers, even in the presence of intervening strong stimuli such as a high concentration of sucrose solution. This also indicated that worker bees do not prefer royal jelly.

On the other hand, the proximity in PER ratios for the mixture and sucrose solution given by 1-day-old bees may be due to the protein affinity of young bees. Protein intake is crucial for the development of the hypopharyngeal glands of 1-day-old workers, and they consume pollen and receive protein-abundant jelly from older nurse bees (Crailsheim, 1990a; Free, 1957; Crailsheim, 1990b). The positive response of 1-day-old workers in this study towards the mixture solution containing royal jelly could then be the typical feeding response of these bees due to the rich protein content of royal jelly (Ramanathan et al., 2018).

Queens exhibited a near-maximal response to both royal jelly and the mixture throughout the trials. Notably, the queens’ response to royal jelly-based stimuli remained consistently high throughout the trials without habituation (Figure 1C). This high and persistent response underscores a highly specialized sensory tuning. Because the queen’s reproductive output is dependent on a royal jelly diet (Fèvre & Dearden, 2024). Queens can be fed nectar and pollen by nurse bees, and they can feed individually on these sources (St Clair et al., 2024). Moreover, when the queens were separated from the court bees, they could consume sugar. However, only sugar as food decreases the queen’s longevity (Haydak, 1970). Additionally, there were only trace amounts of sugar found in the gut of the queens during the winter period (Haydak, 1970). Thus, queen bees are responsive to sucrose, but sugar is not an adequate or preferred food for them.

Our findings on sucrose response are also consistent with the literature. Queen bees responded to sugar in this study, as shown before (Gong et al., 2018). Older bees showed a higher affinity for sugar than younger bees in this study, as shown by higher PER to sugar by foragers, as in other studies (Pankiw & Page, 1999; Guez et al., 2001; Scheiner et al., 2003; Degirmenci et al., 2018). We also found that the 1-day-old workers had more apparent habituation to sucrose than foragers, in line with previous studies (Guez et al., 2001; Scheiner et al., 2003).

The novel finding of our study is that bees have caste and age-related food preferences. High responsiveness to pure royal jelly is exclusive to the queen bee. Forager bees have a high level of fidelity to sugar, and 1-day-old bees do show some preference for royal jelly in a mixture. One practical implication of this finding is the ability to independently manipulate the nutritional status of members of a hive to examine the nutritional ecology and physiology of the hive. Future research may also unravel the molecular and neural basis of these caste-specific feeding preferences. For example, investigating the differential expression of gustatory and olfactory receptors across castes may explain how queens and workers are biochemically tuned to their respective diets. Sensory neurons responsive to protein-rich stimuli like royal jelly remain poorly characterized in queens, and their comparison with carbohydrate-sensitive pathways in foragers could provide insight into the adaptive evolution of nutritional specialization in eusocial insects.

## ACKNOWLEDGMENTS AND FUNDING INFORMATION

This work was supported by the EU grant RoboRoyale [grant number: 964492]; the Middle East Technical University Research Fund [grant number: ADEP-302-2024-11468]; the U.S. National Science Foundation grant [grant number: 2318597].

## DATA AVAILABILITY

The data are available in the Zenodo repository: https://doi.org/10.5281/zenodo.18347610

## AUTHOR CONTRIBUTIONS

BE: Conceptualization, Methodology, Formal analysis, Investigation, Writing - Original Draft, Visualization. SS: Investigation, Resources. OCA: Investigation, Writing – review and editing. AGG: Writing - Original Draft. HA: Methodology, Writing – review and editing AET: Methodology, Resources, Funding acquisition. TG: Conceptualization, Methodology, Formal analysis, Writing - Original Draft. ES: Conceptualization, Writing - Original Draft, Resources, Funding acquisition.

## REFERENCES

Bayram, E. N., Çebi, N., Çelik, S., Gerçek, Y. C., Bayram, S., Tanuğur Samancı, A. E., Sağdıç, O., & Özkök, A. (2021). Turkish royal jelly: amino acid, physicochemical, antioxidant, multi-elemental, antibacterial and fingerprint profiles by analytical techniques combined with chemometrics. Journal of Apicultural Research, 60(5), 751–764. 10.1080/00218839.2021.1889222

Bitterman, M.E., Menzel, R., Fietz, A. & Schäfer, S. (1983) Classical Conditioning of Proboscis Extension in Honeybees (Apis mellifera). Journal of Comparative Psychology, 97, 107. 10.1037/0735-7036.97.2.107

de Groot, A. P. (1952). Amino acid requirements for growth of the honeybee (Apis mellifica L.). Experientia, 8(5), 192–194. 10.1007/BF02173740

Crailsheim, K. (1990a). The protein balance of the honey bee worker. Apidologie, 21(5), 417–429. 10.1051/apido:19900504

Crailsheim, K. (1990b). Protein synthesis in the honeybee (Apis mellifera L.) and trophallactic distribution of jelly among imagos in laboratory experiments. Zoologische Jahrbucher, Allgemeine Zoologie Und Physiologie Der Tiere, 94, 303–312. https://www.cabidigitallibrary.org/doi/full/10.5555/19910230837

Crailsheim, K., Schneider, L. H. W., Hrassnigg, N., Bühlmann, G., Brosch, U., Gmeinbauer, R., & Schöffmann, B. (1992). Pollen consumption and utilization in worker honeybees (Apis mellifera carnica): Dependence on individual age and function. Journal of Insect Physiology, 38(6), 409–419. 10.1016/0022-1910(92)90117-V

Crailsheim, K., & Stolberg, E. (1989). Influence of diet, age and colony condition upon intestinal proteolytic activity and size of the hypopharyngeal glands in the honeybee (Apis mellifera L.). Journal of Insect Physiology, 35(8), 595–602. 10.1016/0022-1910(89)90121-2

de Brito Sanchez, M. G. (2011). Taste perception in honey bees. Chemical Senses, 36(8), 675–692. 10.1093/chemse/bjr040

Değirmenci, L., Thamm, M., & Scheiner, R. (2018). Responses to sugar and sugar receptor gene expression in different social roles of the honeybee (Apis mellifera). Journal of Insect Physiology, 106(May 2017), 65–70. 10.1016/j.jinsphys.2017.09.009

Fahrbach, S. E., Giray, T., & Robinson, G. E. (1995). Volume changes in the mushroom bodies of adult honey bee queens. Neurobiology of Learning and Memory, 63(2), 181–191. 10.1006/nlme.1995.1019

Farina, W. M., & Núñez, J. A. (1991). Trophallaxis in the honeybee, Apis mellifera (L.) as related to the profitability of food sources. Animal Behaviour, 42(3), 389–394. 10.1016/S0003-3472(05)80037-5

Fèvre, D. P., & Dearden, P. K. (2024). Influence of nutrition on honeybee queen egg-laying. Apidologie, 55(4), 1–16. 10.1007/s13592-024-01097-1

Free, J. B. (1957). The transmission of food between worker honeybees. The British Journal of Animal Behaviour, 5(2), 41–47. 10.1016/S0950-5601(57)80023-9

Giray, T., Galindo-Cardona, A., & Oskay, D. (2007). Octopamine influences honey bee foraging preference. Journal of Insect Physiology, 53(7), 691–698. 10.1016/j.jinsphys.2007.03.016

Guez, D., Suchail, S., Gauthier, M., Maleszka, R., & Belzunces, L. P. (2001). Contrasting effects of Imidacloprid on habituation in 7- and 8-day-old honeybees (Apis mellifera). Neurobiology of Learning and Memory, 76(2), 183–191. 10.1006/nlme.2000.3995

Gong, Z., Tan, K., & Nieh, J. C. (2018). First demonstration of olfactory learning and long-term memory in honey bee queens. Journal of Experimental Biology, 221(14). 10.1242/jeb.177303

Haydak, M. H. (1970). Honey bee nutrition. Annual Review of Entomology, 15(1), 143–156. 10.1146/annurev.en.15.010170.001043

Howe, S. R., Dimick, P. S., & Benton, A. W. (1985). Composition of freshly harvested and commercial royal jelly. Journal of Apicultural Research, 24(1), 52–61. 10.1080/00218839.1985.11100649

Kolayli, S., Sahin, H., Can, Z., Yildiz, O., Malkoc, M., & Asadov, A. (2016). A member of complementary medicinal food: Anatolian royal jellies, their chemical compositions, and antioxidant properties. Journal of Evidence-Based Complementary & Alternative Medicine, 21(4), NP43–NP48. 10.1177/2156587215618832

Lindauer, M. (1952). Ein beitrag zur frage der arbeitsteilung im bienenstaat. Zeitschrift Für Vergleichende Physiologie, 34(4), 299–345. 10.1007/BF00298048

Moreno, E., & Arenas, A. (2023). Changes in resource perception throughout the foraging visit contribute to task specialization in the honey bee Apis mellifera. Scientific Reports, 13(1), 1–10. 10.1038/s41598-023-35163-y

Moreno, E., & Arenas, A. (2024). Foraging task specialization in honey bees (Apis mellifera): the contribution of floral rewards to the learning performance of pollen and nectar foragers. Journal of Experimental Biology, 227(13). 10.1242/jeb.246979

Nicholls, E., & Hempel de Ibarra, N. (2013). Pollen elicits proboscis extension but does not reinforce PER learning in honeybees. Insects, 4(4), 542–557. 10.3390/insects4040542

Nicholls, E., Krishna, S., Wright, O., Stabler, D., Krefft, A., Somanathan, H., & Hempel de Ibarra, N. (2019). A matter of taste: the adverse effect of pollen compounds on the preingestive gustatory experience of sugar solutions for honeybees. Journal of Comparative Physiology A: Neuroethology, Sensory, Neural, and Behavioral Physiology, 205(3), 333–346. 10.1007/s00359-019-01347-z

Pain, J., & Maugenet, J. (1966). Physiologiques sur le pollen par les abeilles emmagasin par les abeiles. Les Annales de l’Abeille, 9(3), 209–236. https://hal.science/hal-00890236v1

Pankiw, T., Nelson, M., Page, R. E., & Fondrk, M. K. (2004). The communal crop: Modulation of sucrose response thresholds of pre-foraging honey bees with incoming nectar quality. Behavioral Ecology and Sociobiology, 55(3), 286–292. 10.1007/s00265-003-0714-0

Pankiw, T., & Page, R. E. (1999). The effect of genotype, age, sex, and caste on response thresholds to sucrose and foraging behavior of honey bees (Apis mellifera L.). Journal of Comparative Physiology - A Sensory, Neural, and Behavioral Physiology, 185(2), 207–213. 10.1007/s003590050379

Pankiw, T., & Page Jr, R. E. (2001). Brood pheromone modulates honeybee (Apis mellifera L.) sucrose response thresholds. Behavioral Ecology and Sociobiology, 49(2–3), 206–213. 10.1007/s002650000282

Ramanathan, A. N. K. G., Nair, A. J., & Sugunan, V. S. (2018). A review on royal jelly proteins and peptides. Journal of Functional Foods, 44(December 2017), 255–264. 10.1016/j.jff.2018.03.008

Rankin, C. H., Abrams, T., Barry, R. J., Bhatnagar, S., Clayton, D. F., Colombo, J., Coppola, G., Geyer, M. A., Glanzman, D. L., Marsland, S., McSweeney, F. K., Wilson, D. A., Wu, C. F., & Thompson, R. F. (2009). Habituation revisited: An updated and revised description of the behavioral characteristics of habituation. Neurobiology of Learning and Memory, 92(2), 135–138. 10.1016/j.nlm.2008.09.012

R Core Team (2020). R: a language and environment for statistical computing. https://www.R-project.org/.

Scheiner, R., Barnert, M., & Erber, J. (2003). Variation in water and sucrose responsiveness during the foraging season affects proboscis extension learning in honey bees. Apidologie, 34(1), 67–72. 10.1051/apido:2002050

Scheiner, R., Page, R. E., & Erber, J. (2001). The effects of genotype, foraging role, and sucrose responsiveness on the tactile learning performance of honey bees (Apis mellifera L.). Neurobiology of Learning and Memory, 76(2), 138–150. 10.1006/nlme.2000.3996

Scheiner, R., Page, R. E., & Erber, J. (2004). Sucrose responsiveness and behavioral plasticity in honey bees (Apis mellifera). Apidologie, 35(2), 133–142. 10.1051/apido:2004001

Seeley, T. D. (1982). Adaptive significance of the age polyethism schedule in honeybee colonies. Behavioral Ecology and Sociobiology, 11(4), 287–293. 10.1007/BF00299306

Simcock, N. K., Gray, H. E., & Wright, G. A. (2014). Single amino acids in sucrose rewards modulate feeding and associative learning in the honeybee. Journal of Insect Physiology, 69(C), 41–48. 10.1016/j.jinsphys.2014.05.004

St. Clair, A. L., Dwyer, B., Shapiro, M., & Dolezal, A. G. (2024). Adult honey bee queens consume pollen and nectar. BioRxiv. 10.1101/2024.12.04.626851

