## Supplementary for "Caste-Specific Proboscis Extension Responses in Honey Bees to Sucrose and Royal Jelly Stimuli"

Table S1. Tukey-adjusted pairwise comparisons of stimuli within each bee category from the GLMM. For each category, the table reports all stimulus-by-stimulus contrasts as odds ratios (exponentiated log-odds contrasts), corresponding z-statistics, and Tukey-adjusted p-values. Significance is coded as asterisks, and “ns” indicates not significant.

| **Category** | **Stimulus 1** | **Stimulus 2** | **Odds Ratio** | **z-statistics** | **P value** | **Significance level** |
|---|---|---|---|---|---|---|
| 1-day old | Water | Sucrose | 0.090 | -11.265 | <0.0001 | **** |
| | Water | Royal Jelly | 0.653 | -2.272 | 0.105 | ns |
| | Water | Mixture | 0.323 | -5.885 | <0.0001 | **** |
| | Sucrose | Royal Jelly | 7.236 | 9.452 | <0.0001 | **** |
| | Sucrose | Mixture | 3.587 | 6.158 | <0.0001 | **** |
| | Royal Jelly | Mixture | 0.496 | -3.712 | 0.001 | *** |
| Forager | Water | Sucrose | 0.021 | -14.274 | <0.0001 | **** |
| | Water | Royal Jelly | 1.019 | 0.104 | 1.000 | ns |
| | Water | Mixture | 0.975 | -0.140 | 0.999 | ns |
| | Sucrose | Royal Jelly | 48.098 | 14.344 | <0.0001 | **** |
| | Sucrose | Mixture | 46.051 | 14.192 | <0.0001 | **** |
| | Royal Jelly | Mixture | 0.957 | -0.244 | 0.995 | ns |
| Queen | Water | Sucrose | 0.007 | -10.315 | <0.0001 | **** |
| | Water | Royal Jelly | 0.000 | -8.235 | <0.0001 | **** |
| | Water | Mixture | 0.000 | -8.181 | <0.0001 | **** |
| | Sucrose | Royal Jelly | 0.009 | -4.307 | 0.0001 | **** |
| | Sucrose | Mixture | 0.010 | -4.249 | 0.0001 | **** |
| | Royal Jelly | Mixture | 1.064 | 0.043 | 1.000 | ns |

Table S2. Tukey-adjusted pairwise comparisons of bee categories within each stimulus from the GLMM. For each stimulus, the table reports all stimulus-by-stimulus contrasts as odds ratios (exponentiated log-odds contrasts), corresponding z-statistics, and Tukey-adjusted p-values. Significance is coded as asterisks, and “ns” indicates not significant.

| **Stimulus** | **Category 1** | **Category 2** | **Odds Ratio** | **z-statistics** | **P value** | **Significance level** |
|---|---|---|---|---|---|---|
| Water | 1-day old | Forager | 1.216 | 0.444 | 0.897 | ns |
| | 1-day old | Queen | 10.422 | 3.784 | <0.0001 | **** |
| | Forager | Queen | 8.571 | 3.483 | 0.001 | *** |
| Sucrose | 1-day old | Forager | 0.286 | -2.571 | 0.027 | * |
| | 1-day old | Queen | 0.810 | -0.340 | 0.938 | ns |
| | Forager | Queen | 2.836 | 1.624 | 0.236 | ns |
| Royal Jelly | 1-day old | Forager | 1.898 | 1.457 | 0.312 | ns |
| | 1-day old | Queen | 0.001 | -5.680 | <0.0001 | **** |
| | Forager | Queen | 0.001 | -6.205 | <0.0001 | **** |
| Mixture | 1-day old | Forager | 3.666 | 2.945 | 0.009 | ** |
| | 1-day old | Queen | 0.002 | -5.052 | <0.0001 | **** |
| | Forager | Queen | 0.001 | -6.116 | <0.0001 | **** |

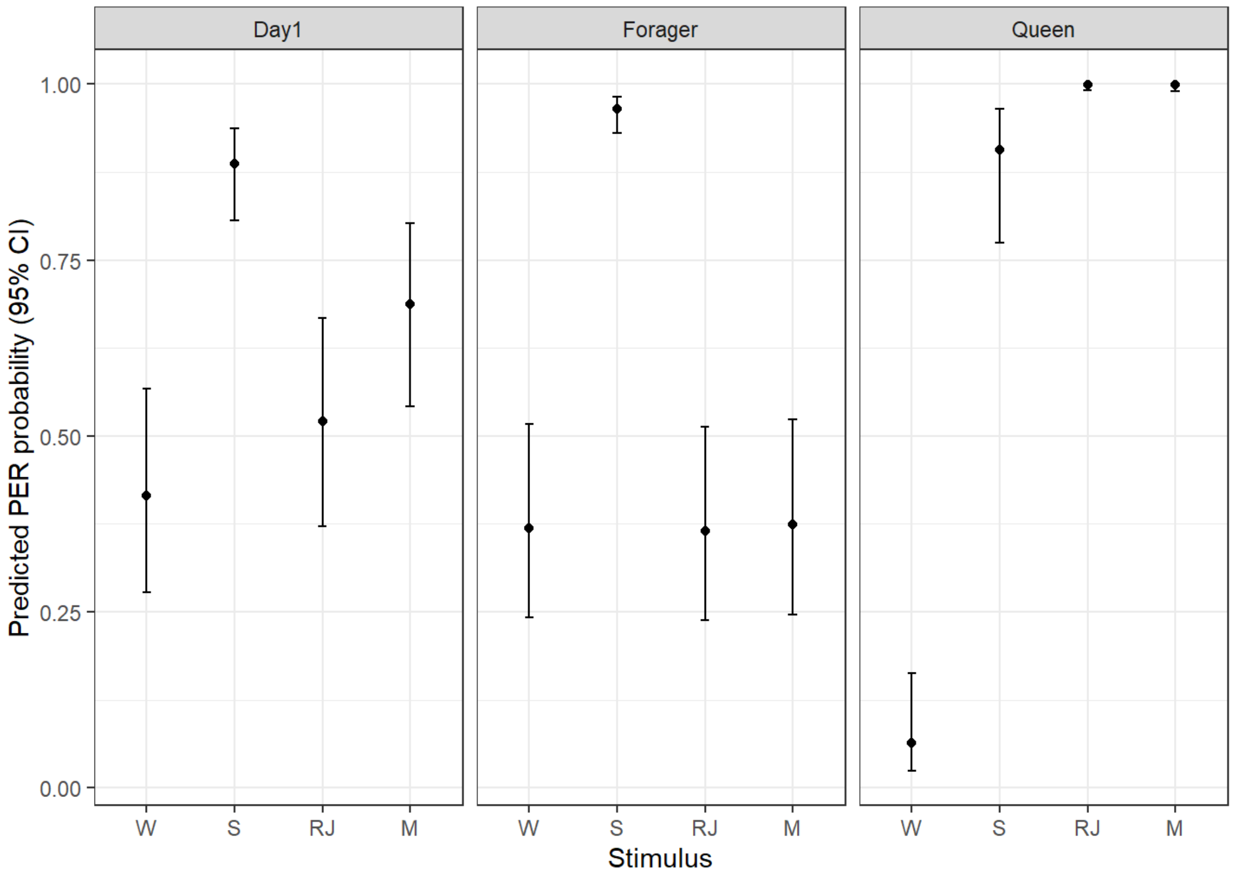


Figure S1. GLMM-estimated PER probabilities by stimulus and bee category. Points show estimated marginal mean PER probability (back-transformed from the logit scale) from the binomial GLMM for each stimulus within each bee category; error bars indicate 95% confidence intervals. S indicates sucrose, W indicates water, RJ indicates royal jelly, and M indicates sucrose–royal jelly mixture.
